# MACROPHAGES COORDINATE IMMUNE RESPONSE TO LASER-INDUCED INJURY VIA EXTRACELLULAR TRAPS

**DOI:** 10.1101/2023.10.16.562553

**Authors:** Federica M. Conedera, Despina Kokona, Martin S. Zinkernagel, Jens V. Stein, Clemens Alt, Volker Enzmann, Charles P. Lin

## Abstract

Macrophages/monocytes, the primary contributors to chronic inflammation in degenerated retinas, orchestrate intricate immune responses. They remain enigmatic in their local coordination and activation mechanisms. Innovations in experimental systems enable real-time exploration of immune cell interactions and temporal dimensions in response. In preclinical mouse models, we use *in vivo* microscopy to unravel how macrophages/monocytes govern microglia and PL responses spatio-temporally.

Our findings underscore the pivotal role of innate immune cells, especially macrophages/monocytes, in regulating retinal repair. The absence of neutrophil and macrophage infiltration aids parenchymal integrity restoration, while their depletion, particularly macrophages/monocytes, impedes vascular recovery. Innate immune cells, when activated, release chromatin and granular proteins, forming extracellular traps (ETs), critical for tissue repair by modulating neutrophil and T-cell responses.

Our investigations demonstrate that pharmacological inhibition of ETosis with Cl-amidine enhances retinal and vascular repair, surpassing the effects of blocking innate immune cell recruitment. Simultaneously, Cl-amidine treatment reshapes the inflammatory response, causing neutrophils, helper, and cytotoxic T-cells to cluster primarily in the superficial capillary plexus, affecting retinal microvasculature perfusion. Our data offer novel insights into innate immunity’s role in responding to retinal damage, potentially informing more effective immunotherapeutic strategies for neurodegenerative diseases.

## INTRODUCTION

Retinal degenerative diseases, such as age-related macular degeneration (AMD), are the leading causes of central vision loss in developed countries. While treatments are available and evolving to manage late-stage symptoms of retinal degeneration (e.g., VEGF treatment), no effective therapies to prevent the pathogenesis toward late-stage degeneration exist (1). This is partly due to the complex and multifaceted disease process involving various cellular abnormalities triggered by local inflammation and peripheral leukocytes (PL) influx into the retina (2). In the central nervous system (CNS), including the retina, the local immune response is provided by microglia, which act as neuropathology sensors by removing debris and toxic substances by phagocytosis (3). During degeneration, they react promptly and adopt distinct phenotypes: anti- or pro-inflammatory state. The pro-inflammatory microglia fuel the inflammatory process, while the anti-inflammatory phenotype is associated with improved phagocytic function (4). Simultaneously, the blood-retinal barrier (BRB), which compartmentalizes blood and retinal parenchyma, alters its permeability characteristics, and the transcellular transport of PL increases (5). Thus, innate and adaptive immune cells are recruited from the circulation to the degenerating area, where they can interplay with microglia (6–8).

An effective inflammatory response upon retinal damage requires the coordinated contribution of the local and infiltrated immune cells. Resident microglia and recruited macrophages are essential for regulating tissue remodeling and repair, and for re-establishing retinal homeostasis. These professional phagocytes also play critical roles in initiating inflammation and orchestrating its resolution (9). However, how phagocytic cells coordinate the immune response to retina damage remains unknown.

Recent investigations have demonstrated that microglia and macrophages can produce extracellular traps [ETs; (10, 11)], which consist of strands of DNA studded with histones and cellular proteins (12). ETs can be formed in response to various stimuli, including pro-inflammatory cytokines, which are also involved in the pathogenesis of several ocular diseases (13). During retinal degeneration, persistent ETosis represents a source of self-antigens that boost the inflammatory process, leading to tissue injury (10). Although there is no current data on ETosis related to AMD, recent evidence showed the existence of ETs in eye tissues, specifically the vitreous body and retina of mice with degenerative retinopathy (14). Furthermore, anti-VEGF therapy reduced ET release during retina degeneration (15).

Here, we used a mouse model of retinal injury generated by laser to define the function of ETs during retinal degeneration. For this, we combined minimally invasive techniques with *in vivo* imaging, pharmacological treatments, and knockout mice to determine the role of phagocytes and how they modulate the local inflammatory response through ETs. Dissecting how this happens is crucial for establishing new and improved immunotherapeutic approaches that can guide a specific maladaptive immune response to a favorable wound healing in the retina.

## METHODS

### Animals

All experimental protocols were approved by the Institutional Animal Care and Use Committee of Massachusetts General Hospital. Male and female 12 weeks old C57Bl/6J mice were purchased from Jackson Laboratories (Bar Harbor, ME). Homozygous LysM^GFP^ mice were obtained from Dr. T. Graf (Center for Genomic Regulation, Barcelona, Spain) and outbred to C57Bl/6J mice. LysM^GFP^ animals express a green fluorescent protein (GFP^+^) in neutrophils and macrophages/monocytes (16). Homozygous Cx3cr1^GFP^Ccr2^RFP^ mice [B6.129(Cg)-Cx3cr1tm1Litt Ccr2tm2.1Ifc/JernJ] were acquired from Jackson Laboratory (Strain #032127). To obtain heterozygotes, these mice were bred with C57Bl/6J mice. Cx3cr1^GFP^Ccr2^RFP^ mice express green fluorescent protein (GFP^+^) specifically in microglia and monocytes/macrophages (17–19). Additionally, they exhibit red fluorescent protein (RFP^+^) expression in peripheral leukocytes (PL; 19, 20). The subset of PL that are Ccr2^+^ consists of monocytes/macrophages, T-cells, and neutrophils (19, 21, 22, 23, 24). Monocytes/macrophages can express both Cx3cr1 and Ccr2 (RFP^+^GFP^+^), while microglia are Cx3cr1^+^Ccr2^-^ and therefore only GFP^+^ (25). They were used to assess *in vivo* the inflammatory response to retinal injury. Though sex differences in immune reactivity were previously reported (20), no differences were detected during intravital experiments (data not shown); thus, both male and female animals were used, and the results combined. Mice having a neomycin cassette replacing exons 3 and 4 of the colony-stimulating factor 2 (granulocyte-macrophage) (Csf2) gene (GM-CSF; B6.129S-Csf2tm1Mlg/J, Strain #026812) were purchased from Jackson Laboratory (Bar Harbor, ME). GM-CSF stimulates the production of granulocytes (neutrophils, eosinophils, and basophils) and monocytes; thus, these mice have impaired innate immunity (GM-CSF1 KO). Homozygous mice for fractalkine receptor (Cx3cr1^gfp/gfp^; B6.129P-Cx3cr1tm1Litt/J) were also obtained from Jackson Laboratory. Part of Cx3cr1^gfp/gfp^ mice was maintained either as homozygotes (Strain #005582) to investigate the dysfunction of microglia-macrophages during injury response or as heterozygous for studying their behavior *in vivo* (21, 22). Csf1r^GFP^ mice were acquired from Jackson Laboratory (Strain #005070) and express enhanced GFP in phagocytic cells of the retina, such as macrophages and microglia. All animals were housed in designated animal holding facilities observing a standard twelve-hour day/night cycle. Standard rodent chow and water were provided *ad libitum*.

### Intravital imaging of the murine retina

Our scanning laser ophthalmoscope (SLO) was customized for multi-color confocal imaging of the murine retina (23, 24), starting from the microscope described by Veilleux et al. (25).

A 638 nm diode laser (Micro Laser Systems, Inc. Garden Grove, CA, USA) detects reflected light and excites AlexaFluor647 (AF647). A two-color laser (Dual Calypso, Cobolt AB, Vretenvägen, Sweden) excites 532 nm to detect GFP/fluorescein and RFP signals at 491 and 532 nm, respectively. A spinning polygon scanner (Lincoln Laser Corp., Phoenix, AZ, USA) and a galvanometric mirror (GSI Lumonics, Billerica, MA, USA) raster scan a field of view of 425 to 575 µm on the retina at 30 frames per second. The laser power incident on the cornea is typically 0.6 mW for the 638 nm diode laser, 0.5 mW for the 532 nm laser, and 0.5 mW laser power for the 491 nm laser. Reflectance images were obtained by splitting the backscattered from the vertically polarized incident light. This was achieved by means of a quarter-wave plate and a polarizing beam splitter cube that generated rotated linear polarization. Light reflected from the retina is horizontally polarized after the double-pass through the quarter-wave plate and is reflected by the polarizing beam splitter, focused through a confocal pinhole (diameter 25 µm ≈ 1.25 airy disc sizes) and detected by a photomultiplier tube (PMT; R3896, Hamamatsu, Japan). Fluorescence from the retina was conducted into a fluorescence detection arm via a dichroic beam splitter (Di03-R405/488/532/638, Semrock, Rochester, NY, USA), separated into three channels by 560 nm and 650 long-pass dichroic beam splitters, and detected through a 650 nm long-pass filter (AF647, Evan Blue), 525/50 (GFP), and 593/40 (RFP) band-pass filters (all Semrock). All fluorescence channels identify light through a confocal pinhole (diameter 50 µm ≈ 2.5 airy disc sizes) with a PMT (R3896, Hamamatsu, Japan).

Mice were placed in a heated holder integrated with a nose cone for inhalation anesthesia (1%–2% isoflurane in oxygen), and their pupils dilated with tropicamide 1% ophthalmic solution (Bausch & Lomb Inc., Tampa, FL, USA). A contact lens (diameter 2.5 mm, base curvature 1.65 mm, power: þ12D, material PMMA; Unicon Corporation, Osaka, Japan) was applied on the mydriatic eye, and a drop of GenTeal eye gel (Alcon, Fort Worth, TX, USA) prevented dehydration of the cornea. Once positioned in the SLO such that the field of view was focused around the optic nerve head (ONH), 10 video-rate movies of fluorescence and reflectance channels were streamed to disk, each approximately 30 seconds in length. *In vivo* images were recorded before injury (baseline) and at days 0, 1, 4, 7, 10 and 14 after laser damage. At the end of the procedure, mice were returned to their cages where they became fully ambulatory within ∼10 minutes.

### Retinal laser injury

We generated focal damage through SLO image guidance using a coagulator embedded in the imaging system. The coagulator includes a high-power continuous-wave laser (Ventus, Laser Quantum, Cheshire, UK) and an acousto-optic modulator (AOM; TEM-85-1-.532, Brimrose, Baltimore, MD, USA) that allows pulses to be chopped from the laser emission. A small mirror reflected the 0-order beam of the AOM into a beam dump; the first-order beam was directed onto a tip-tilt-scanner. The scanner was located in a plane conjugate with the mouse eye pupil through a telecentric relay system of two 75 mm focal length lenses. The scanner served to position the coagulator spot onto a retinal parenchyma. The coagulation beam was combined with the SLO excitation lasers by a dichroic beam splitter (DiO1-R532, Semrock, Rochester, NY). This beam splitter allows placing 532 nm laser pulses to the retinal parenchyma under the guidance of simultaneous real-time imaging by the SLO using the 638 nm laser.

Anesthetized mice received a single laser burn in the nasal region of the retina at a distance of at least one lesion diameter from the ONH. The burn has a diameter of 100 µm in diameter and was performed with a 25 ms pulse to prevent any potential collateral damage to the surrounding tissue adjacent to the laser spot.

### Fluorescein angiography (FA)

For visualizing the vascular plexus, we injected intravenously 50 μL of 0.01% fluorescein (AK-Fluor® 10%; Akorn Inc., Lake Forest, IL, USA). We performed dye injection while the mouse was on stage, because of the high permeability of the BRB during degeneration. Thus, the early dynamics of dye leakage directly after injection can be analyzed *in vivo*. We recorded videos simultaneously in the reflectance and fluorescence channels before (baseline), after laser treatment (day 0) and on days 1, 4, 7, 10 and 14 post injury.

### Labeling of cells for *in vivo* imaging

C57Bl/6J or LysM^GFP^ mice were injected intravenously with 5 µg of AF647 conjugated to anti-Ly6G antibody (127610; Biolegend, San Diego, CA USA) 3 hours before imaging. Antibodies were re-injected on days 1, 3, 7, 10, and 14 for longitudinal imaging of the innate immune response upon injury. Furthermore, C57Bl/6J mice were injected intravenously with 5 µg of AF488 conjugated to anti-CD4 antibody (100529; Biolegend) and 5 µg AlexaFluor647 conjugated to anti-CD8 antibody (100724; Biolegend) 40 minutes before imaging. Antibodies were re-injected on days 1, 3, 7, 10, and 14 post-injuries for longitudinal imaging. Repeated doses of antibodies at a low concentration avoid cell depletion, such that cells could be quantified for 8-10 hours. To visualize ETs, LysM^GFP^ and Cx3cr1^GFP^ mice received 5 μL i.v. Sytox Deep Red (S11380, Invitrogen, Waltham, MA, USA) 30 min before imaging. Intravenously injected tracers were detected by acquiring still images.

A time-lapse image sequence was obtained by taking one picture every 30 seconds over 5 minutes. The time-lapse stack was aligned to rectify any motion artifacts that might have occurred during the imaging process. Then, an average image was generated from the aligned stack to reduce image noise. The presented images were contrast stretched to the same white and black levels for display purposes. All image processing was done using ImageJ.

### Pharmacological treatments

A stock solution of PAD4 inhibitor (Cl-amidine; 506282, Merck, Darmstadt, Germany) was dissolved in dimethyl sulfoxide (DMSO) (Sigma-Aldrich). The stock solution was then dissolved in saline (5% v/v) and directly injected i.p. at 10 mg/kg 24 h after laser injury and every day until the end of experiments. The control group received an injection of the vehicle (saline containing 5% DMSO).

### Transmission electron microscopy (TEM)

Eyes were enucleated and fixed with Karnovsky’s fixative (1% paraformaldehyde (PFA), 3% sodium cacodylate–HCl, and 3% glutaraldehyde; Sigma-Aldrich, Buchs, Switzerland) for 24 h. After washing three times in TEM buffer (2.5% glutaraldehyde and 0.1 M sodium cacodylate–HCl), the eyes were post-fixed in 4% osmium tetroxide in cacodylate buffer for 15 min. Samples were dehydrated through a graded series of acetone, washed with a resin/1,2-propylene oxide mixture (Merck, Darmstadt, Germany), and embedded in epoxy-based resin. Ultrathin sections (80 nm) were cut with an ultramicrotome Ultracut E (Reichert Microscope Services, Depew, NY, USA) equipped with a 45° diamond knife (Diatome, Biel, Switzerland). Sections were placed onto copper grids (G100H-C3; Science Services, Munich, Germany) and counterstained with 4% aqueous uranyl acetate and 0.1% Reynolds’ lead citrate (Science Services). TEM analyses were performed on a CM 12 electron microscope (Philips Applied Technologies, Eindhoven, The Netherlands).

### Retinal dissociation, sorting, and RNA-Seq library production

Both retinas of three Csf1r^GFP^ mice were dissected at different timepoints (Day 1, 3, and 7) and incubated with papain (Worthington Biochemical, Freehold, NJ, USA) for 15 min as previously described (26). After dissociation, cell suspension in HBSS with 0.4% BSA (ThermoFisher Scientific, Basel, Switzerland) and DNase I (200 U/ml; Sigma-Aldrich) was filtered with a 35 μm cell strainer. Hoechst 33342 Ready Flow™ Reagent (ThermoFisher Scientific) was added as DNA dye for cell viability. Cells from Csf1r^GFP^ negative littermates were used to determine background fluorescence levels. 100 cells/μl were sorted from Csf1r^GFP^ mice using Moflo Astrias (Beckman-Coulter, Nyon, Switzerland) into 4 μl Buffer TCL (1,031,576; Qiagen, Venlo, Netherlands) plus 1% 2-mercaptoethanol (Sigma-Aldrich). After cell sorting, all samples were processed using the published Smart-seq2 protocol to generate the cDNA libraries (27). The libraries were sequenced in an Illumina HiSeq4000 (Illumina, San Diego, CA, USA) with a depth of around 20 Mio reads per sample.

### RNA-seq analysis

The raw reads were first cleaned by removing adapter sequences, trimming low quality ends, and filtering reads with low quality (phred quality < 20) using Trimmomatic (Version 0.36). The read alignment was done with STAR (v2.6.0c). As reference the Ensembl murine genome build GRCm38.p5 with the gene annotations downloaded on 2018-02-26 from Ensembl (release 91) were used. The STAR alignment options were “--outFilterType BySJout --outFilterMatchNmin 30 --outFilterMismatchNmax 10 --outFilterMismatchNoverLmax 0.05 --alignSJDBoverhangMin 1 --alignSJoverhangMin 8 --alignIntronMax 1000000 --alignMatesGapMax 1000000 --outFilterMultimapNmax 50”. Gene expression values were computed with the function featureCounts from the R package Rsubread (v1.26.0). The options for feature counts were: - min mapping quality 10 - min feature overlap 10 bp - count multi-mapping reads - count only primary alignments - count reads also if they overlap multiple genes. To detect differentially expressed genes, we applied a count based negative binomial model implemented in the software package DESeq2 (R version: 3.5.0, DESeq2 version: 1.20.0). The differential expression was assessed using an exact test adapted for over-dispersed data. Genes showing altered expression with an adjusted p-value < 0.05 (Benjamini and Hochberg method) were considered differentially expressed. Heatmaps were generated for selected subsets of genes in R v. 3.5.1 using the heatmap.2 function from package gplots v. 3.0.1. The data displayed the log2 fold-changes between two experimental groups.

### Quantification and statistical analysis

We determined damaged area of the laser lesion in reflectance. The damaged site was identified as a hyper-reflective tissue (10% higher intensity compared to healthy parenchyma) bordered by a hypo-reflective circle (10% lower intensity compared to healthy parenchyma). We manually determined the number of positive cells, the injury and leakage area from *in vivo* experiments using ImageJ/Fiji (28). The segmentation process for identifying Cx3cr1^+^, Ccr2^+^, CD4^+^, and CD8^+^ cells from pictures obtained by *in vivo* imaging was done as previously reported (29). Representative figures show the image processing for differentiating LysM^GFP^ and Ly6G^+^ cells from pictures obtained by *in vivo* imaging (**Fig. S1**).

Microglia polarization toward the damage site was defined based on the calculation of the polarization coefficient P as follows: P = [AVG (Dp - Ds)]/Kd, where Dp = distance of process tip from the center of the damage, Ds = distance of soma from the center of the damage, and Kd = average diameter of microglia. We measured P of each microglia found in the field of view (≈425 µm) and then averaged our measurements per animal. Each time point (days 1, 4, 7, 10, 14) was normalized to day 0 (after injury).

We employed the VasoMetrics macro in ImageJ/Fiji to quantify the capillary plugs, ensuring minimized measurement bias (26, 27). The process involved the following steps: initially, the user drew a line that bisected the targeted vessel segment. Subsequently, the macro generated perpendicular cross-lines, spaced 3 µm apart, to capture the fluorescence intensity across the vessel’s width. By calculating the full width at half-maximum of the intensity profile at equidistant locations, we obtained multiple diameter values for each segment. These values were then averaged and converted to micrometer units, resulting in a single diameter value per vessel segment. Our analysis classified a capillary as non-flowing if it exhibited filling defects in the FA. Additionally, we excluded counts of transitory capillary stalling, as they were attributed to temporary flow slowdowns lasting less than 5 seconds, typically caused by the passage of large immune cells through a capillary.

The Spearman’s correlation matrix was generated using the corrplot function from the gplots package (version 3.0.1) in R v. 3.5.1. All measurements were conducted blinded to the treatment groups, and normality tests were carried out prior to the statistical analyses, when suitable. Parametric analyses (t-test, one-way ANOVA with Holm-Sidak or Bonferroni post-hoc test for multiple comparisons) were performed and reported as means when the data met a normal distribution. In cases where normality was not met, non-parametric analyses (Mann–Whitney test, Spearman rank-sum correlation) were used, and median values were reported. For longitudinal measurements, linear mixed-effects modeling was utilized, and the specific test used is described in the text and figure legends. Statistical significance was considered at a p-value ≤ 0.05. All statistical analyses were conducted using GraphPad Prism 10 (GraphPad Software, Boston, MA, USA).

## RESULTS

### Characterization of the innate immune response to laser-induced injury and its effect on tissue repair in the retina

In our study, we employed a scanning laser ophthalmoscope (SLO) to investigate *in vivo* the innate immune response to focal injury in the retina. We specifically focused on two sub-populations of innate immune cells: macrophages/monocytes (LysM^GFP^) and neutrophils (LysM^GFP^/Ly6G^+^). We analyzed the recruitment of these labeled cells at the injury site, particularly by the nerve fiber layer (NFL; superficial) and the photoreceptor layer (PR; deep).

Our findings revealed that innate immune cells were recruited close to the damaged area on both days 1 and 7. This recruitment was observed in the inner retina near the NFL and the outer retina adjacent to the photoreceptors (PR; **Fig. 1 a-e**). Specifically, LysM^GFP^ cells formed clusters above the damage site on days 1 and 7, with a reduction in their response on day 4 (**Fig. 1 b-d**). Conversely, LysM^GFP^/Ly6G^+^ cells predominantly migrated towards the injured site in the proximity of the NFL on day 1, and only a few positive cells were still visible from day 4 onwards until day 10 (**Fig. 1 b-d**). Furthermore, we observed that LysM^GFP^ cells were primarily present around the injured PR on days 1, while LysM^GFP^/Ly6G^+^ cells appeared near the damaged outer retina on days 1 and 7. Similar to the LysM^GFP^ cells, the response of LysM^GFP^/Ly6G^+^ cells decreased on day 4 (**Fig. 1 b-c, e**).

**Fig. 1:**
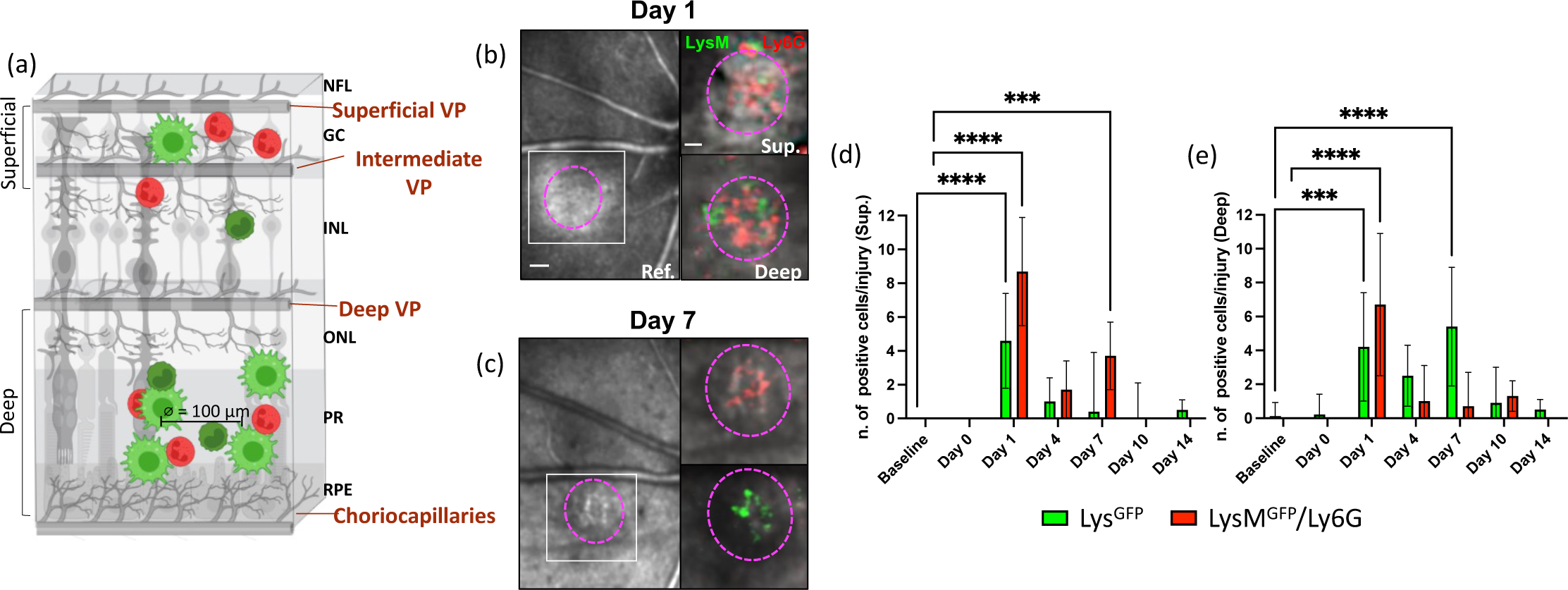
Innate immune response to laser-induced damage in the murine retina. (a) Schematic of the inflammatory response in the retina upon laser-induced injury. (a) The damage is symbolized by a gap in the RPE and PR, where innate immune cells are recruited. We quantified the cell density, discriminating between superficial vasculature nearby the NFL (superficial and intermediate vascular plexus) and the deep vasculature nearby PR and RPE (deep vascular network and choriocapillaris). NFL, nerve fiber layer; VP, vascular plexuses; GC, ganglion cell layer; INL, inner nuclear layer; ONL, outer nuclear layer; PR, photoreceptor layer; RPE, retinal pigment epithelium. (b-c) Tracking of macrophages/monocytes (LysM^GFP^) and neutrophils (LysM^GFP^/Ly6G^+^) in the same eye of a LysM^GFP^ mouse within the damaged area. Magenta dashes outline retinal injury in reflectance (Ref.), while inserts correspond to the region delimited by a white box in reflectance. Inserts show the recruitment of innate immune cells in the injured site on days 1 and 7 by the NLF (Sup.) and the PR (Deep). Scale bar is 100 μm in reflectance images and 50 μm in the inserts. (d) Quantification of the number of macrophages/monocytes and neutrophils per lesion in the proximity of the superficial vasculature at baseline, days 0 (after injury), 1, 4, 7, 10, and 14. Significant differences (***p<0.001 and ****p<0.0001) between baseline and the different time points were determined by using a post-hoc Bonferroni one-way ANOVA test (n=8) (e) Quantification of the number of macrophages/monocytes and neutrophils per lesion in the proximity of the deep vasculature network at baseline and at different time points after laser (Days 0, 1, 4, 7, 10 and 14). Significant differences (***p<0.001 and ****p<0.0001) between baseline and different time points were determined using a post-hoc Bonferroni one-way ANOVA test (n=8).

Overall, our results provide *in vivo* evidence of the recruitment and spatial distribution of innate immune cell populations in response to retinal injury, highlighting distinct patterns of cellular migration and temporal changes in their responses. To examine how innate immunity affected retinal repair, we compared the extent of the injury to the retinal parenchyma and the vascular supply upon injury between untreated and genetically modified mice with altered innate immune responses (**Fig. 2a-d**). We utilized GM-CSF1 KO mice, which are unable to generate neutrophils and macrophages from bone marrow progenitors (30, 31), and Cx3cr1^gfp/gfp^ mice, which exhibit impaired recruitment of macrophages/monocytes (32). During the initial 24 hours, we observed the damaged site characterized by a hypo-reflective signal encompassed by a hyper-reflective circle across all groups (**Fig. 2a**). While the untreated retinas displayed visible injury throughout the experiment, both GM-CSF1 KO and Cx3cr1^gfp/gfp^ mice exhibited a significant reduction in the extent of the damage starting from day 1 (**Fig. 2a-b**). Notably, the injury size showed a 5-fold reduction in GM-CSF1 KO animals compared to untreated mice, and it was even smaller in Cx3cr1^gfp/gfp^ mice on day 4 (**Fig. 2b**). These findings suggest a crucial role for innate immunity in retinal repair, as inhibition of macrophage/monocyte recruitment led to the accelerated recovery of the retinal parenchyma compared to broader depletion of neutrophils and macrophages.

**Fig. 2.**
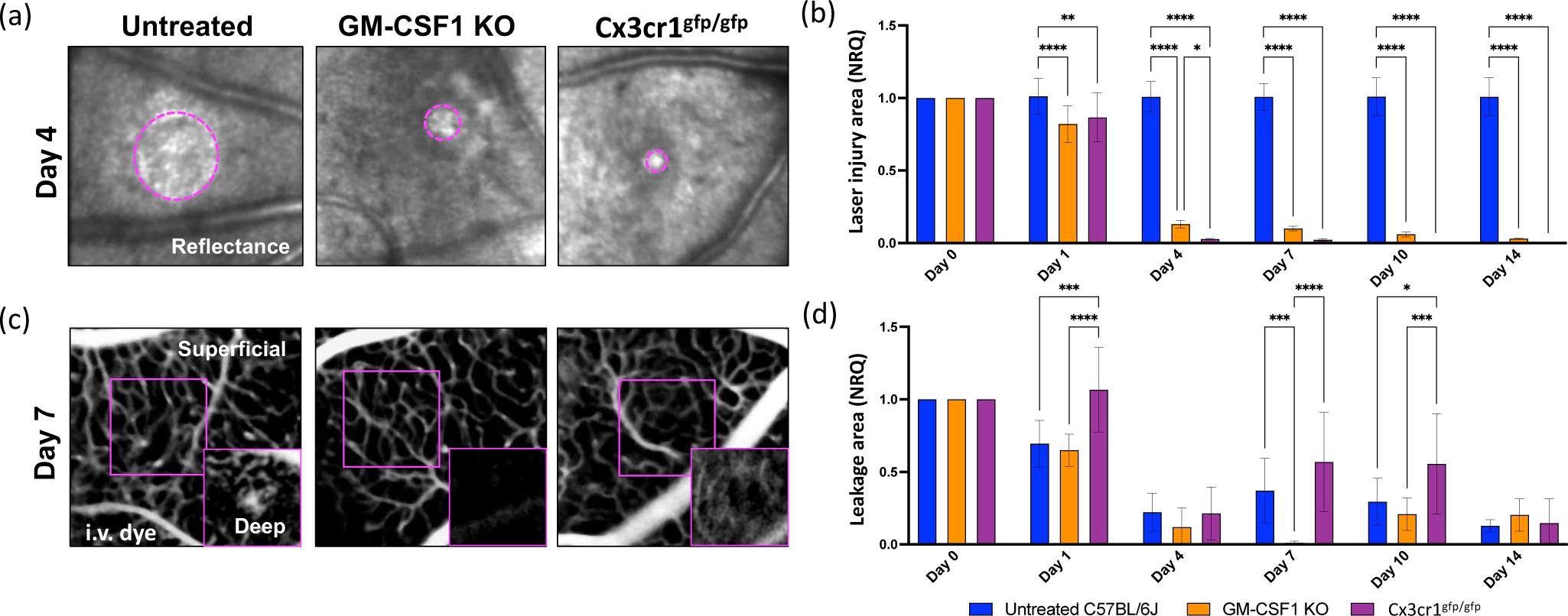
Impact of innate immune response on the recovery of retinal parenchyma and barrier function after laser-induced injury. (a-b) Kinetics of retinal injury detected as back-scattered light (Reflectance) in untreated, GM-CSF1 KO and Cx3cr1^gfp/gfp^ mice. The damage was identified as a hypo-reflective circle (≥10% lower intensity compared to healthy parenchyma) surrounding a hyper-reflective core (≥10% higher intensity compared to healthy parenchyma). (a) Images show a reduction of reflective area in the damaged site (delimited by magenta dashes) of GM-CSF1 KO and Cx3cr1^gfp/gfp^ retinas on day 1 compared to untreated retinas. (b) Quantification of the damaged area of untreated, GM-CSF1 KO and Cx3cr1^gfp/gfp^ retinas after injury (day 0) and on days 1, 4, 7, 10, and 14). Significant differences (*p<0.1, **p<0.01, and ****p<0.0001) between untreated, GM-CSF1 KO and Cx3cr1^gfp/gfp^ mice were determined by using a post-hoc Bonferroni one-way ANOVA test (n=8). (c) Representative angiographs of untreated, GM-CSF1 KO and Cx3cr1^gfp/gfp^ eyes on day 7. (d) Quantification of leakage deep in the retina after injury (day 0) and on days 1, 4, 7, 10, and 14. Significant differences (*p<0.1, ***p<0.001, and ****p<0.0001) between untreated, GM-CSF1 KO and Cx3cr1^gfp/gfp^ mice were determined using a post-hoc Bonferroni one-way ANOVA test (n=8). Day 0 was chosen as the calibrator [NRQ (normalized relative quantification) = 1]. Field of view is ≈425 µm.

Concurrently, we analyzed the integrity of the retinal vasculature during injury response by recording *in vivo* the dynamics of fluorescein leakage immediately after injection. We also discerned whether fluorescein leaked from the superficial vasculature in the proximity of the NFL (superficial and intermediate vascular plexus) or the deep vascular network in the proximity of the PR (**Fig. 2 c-d**). The deep vascular network adjacent to the RPE and PR was compromised until the last time point investigated in all groups, while the superficial vasculature remained intact (**Fig. 2c**). Following laser application, we observed a similar reduction in fluorescein leakage between untreated and GM-CSF1 KO mice on day 1 (**Fig. 2d**). Conversely, Cx3cr1^gfp/gfp^ mice, which exhibited impaired macrophage/monocyte recruitment, displayed no statistically significant difference in sub-retinal fluid accumulation compared to the baseline (**Fig. 2d**). While vascular leakage was comparable among all groups by day 4, both untreated and Cx3cr1^gfp/gfp^ mice exhibited a more pronounced leakage between days 7 and 10 (**Fig. 2c-d**). Interestingly, minimal fluorescein leakage was observed only in the angiograms of GM-CSF1 KO mice on day 7 (**Fig. 2c-d**). On the other hand, the deep vascular network remained compromised without neutrophils and macrophages, as evidenced by the fluorescein signal detected on days 10 and 14 (**Fig. 2d**). These findings strongly indicate the critical role of innate immunity in tissue repair. The absence of neutrophil and macrophage infiltration into the retina contributes to restoring parenchymal integrity. In contrast, depletion of neutrophils and macrophages, particularly macrophages/monocytes, may exacerbate vascular damage.

Our data reinforce the significance of innate immune cells in the repair process, with their absence leading to the rescue of retinal tissue integrity. Conversely, an altered innate immunity may exacerbate vascular damage, underscoring the delicate balance of their involvement in the healing response.

Our data were endorsed by Spearman’s correlations of injury area or dye leakage with macrophages/monocytes and neutrophils in untreated mice (**Fig. S2**). We showed an association only between macrophages/monocytes clustering in the injury and the extent of the injury/vascular damage. No correlation was found with neutrophils during injury response.

### Laser-induced injury induces macrophage ETosis in the retina

Innate immune cells are known to release chromatin and granular proteins forming DNA traps, which are called ETs (33). This process is well characterized in neutrophils, but also occurs in other innate immune cells such as macrophages/monocytes and microglia (10, 12, 34).

Using TEM, we investigated the morphological alterations in phagocytic cells (macrophages and microglia) during the injury response (**Fig. 3 a-e**). Contrary to our expectations, neutrophils displayed a multilobed nucleus, and numerous granules were visible within the cytoplasm after injury. These granules exhibited varying electron densities, some appearing dark (electron-dense) and others lighter (electron-lucent). The cell membrane, forming a thin and continuous boundary, separated the cytoplasm from the extracellular environment (**Fig. S3a**). These fundamental characteristics remain recognizable in neutrophils not undergoing ETosis. On the other hand, phagocytic cells initially experienced a dilation between their inner and outer nuclear membranes, commonly referred to as blebbing (**Fig. 3 a**). Within this separation of membranes, we observed DNA strands with bound nucleosomes in a repeated array, exhibiting an ultrastructural appearance resembling “beads on a string” (**Fig. 3 a**). This characteristic appearance of DNA strands with bound nucleosomes has been previously reported in the literature (35). Between days 4 and 7, we detected vesicles within the cytoplasm of cells, enclosed by ETs, where strands of DNA were also present (**Fig. 3 b-c**). Furthermore, phagocytic cells released dense DNA filaments into the extracellular space (**Fig. 3 c**).

**Fig. 3:**
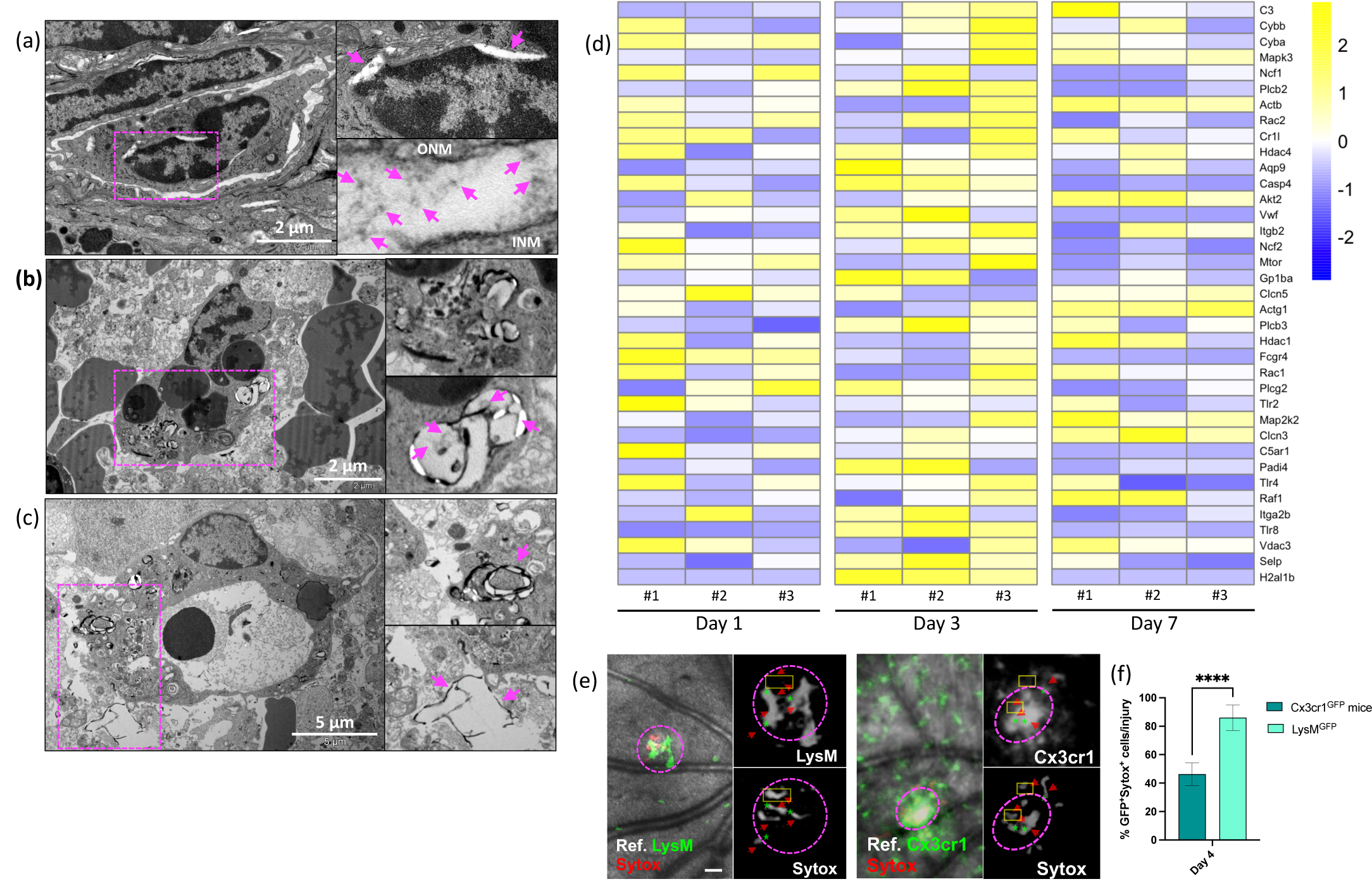
Macrophages produce ETs during injury response in the retina. (a) Transmission electron microscopy of ET formation showing nuclear envelope alterations in phagocytic cells on day 1. Close-up of separation of the inner nuclear membrane (INM) from the outer nuclear membrane (ONM) and small DNA strands within the lumen between the INM and ONM. (b) Images show a phagocytic cell with DNA containing vesicles in the cytoplasm on day 4. Enlarged view of vesicle containing DNA strands. (c) Transmission electron microscopy of phagocytic cells releasing ETs on day 7. Close-up of dense DNA filaments released into the extracellular space. (d) Heatmaps of differentially expressed NETs-related genes in Csfr1GFP cells. Genes were selected from KEGG pathways (mmu04613). Data are expressed as fold-changes between different time points (days 1, 3, and 7) compared to negative controls (Csfr1^EGFP^ cells from uninjured retinas). (e) Representative in vivo images of macrophages of LysMGFP mice (left) or Cx3cr1+ cells (green, right) releasing extracellular DNA (red) on day 4. Extracellular DNA was labeled with intravenous injection of SYTOX Red. Arrows indicate extracellular DNA fibers, and stars are placed on the cell bodies. (f) Percentage of Cx3cr1^+^/SYTOX^+^ and LysM^+^/SYTOX^+^ per lesion 4 days after injury. Significant differences (****p<0.0001) between baseline and the different time points were determined by using a two-tailed Mann-Whitney test analysis (n=8). Field of view is approximately ≈575 µm.

To validate our TEM data, we conducted a transcriptome analysis of phagocytic cells expressing the colony-stimulating factor 1 receptor (Csf1r) before and after injury at specified time points (days 1, 3, and 7). Using RNAseq from Csf1r^+^ cells, we identified 38 genes related to ETosis with significant fold-changes (p-value < 0.05). Notably, most of these genes showed upregulation at day 3 post-injury. Among the significantly upregulated transcripts, we observed cytochrome b (*Cyba* and *Cybb*), Rac family small GTPase 2 (*Rac2*), integrin subunit beta 2 (*Itgb2*), and neutrophil cytosolic factor 2 (*Ncf2*). These genes have previously been shown to directly regulate neutrophil extracellular traps (NET) formation (10, 36–39). We also detected an upregulation of *Casp4* and *Vwf*, both of which are involved in NET formation, facilitating nuclear expansion, lysis, and NET release (10, 40). Furthermore, our analysis revealed increased expression of histone deacetylases (Hdac1 and 4), which are known to play a crucial role in NET formation by allowing peptidyl-arginine deiminase 4 (*Pad4*) mediated histone citrullination, initiating chromatin de-condensation (41–43). As expected, Pad4 itself was consequently upregulated as well. Unlike other critical genes for initiating ET formation, such as *Mpo* and *Ela2* which were instead downregulated compared to baseline levels (**Fig. S4**).

To confirm if either microglia or macrophages are capable of ETosis, we identified ET formation using *in vivo* confocal imaging of a DNA-binding SYTOX dye. We then quantified if microglia (Cx3cr1^+^) and macrophages (LysM^GFP^) released ETs upon laser-induced injury in the retina in homozygous Cx3cr1^GFP^ and LysM^GFP^ mice, respectively (**Fig. 3 d-e, S3 b**). This analysis revealed that Sytox^+^ extracellular DNA fibers were not observed at the baseline, whereas these fibers were abundantly found in the damaged site (**Fig. 3 d**). Since SYTOX is also used to detect apoptosis, its signal was widely diffused within the injured area on day 1. Thus, SYTOX did not co-localize only with the microglia or macrophages (data not shown). However, starting from day 4, SYTOX preferentially localized with LysM^GFP^ cells instead of Cx3cr1^+^ cells (**Fig. 3 e**), suggesting that macrophages produce ETs.

We have provided evidence that macrophages upregulate primarily PAD4 between the genes responsible for initiating chromatin decondensation. This process makes chromatin accessible for release from macrophages, enabling the formation of ETs (44). Therefore, we investigated the role of ETs during retinal injury response and pharmacologically inhibited PAD4, using Cl-amidine (45). We compared the extent of the injured retinal parenchyma and the fluorescein leak upon injury between GM-CSF1 KO, Cx3cr1^gfp/gfp^, PAD4-treated and untreated mice (**Fig. 4 a-d**). In GM-CSF1 KO, Cx3cr1^gfp/gfp^, and PAD4-treated mice, the damaged site showed a similar reduction within 24 hours (**Fig. 4 b**). Starting from day 4, no significant hypo- or hyper-reflectivity was observed in the absence of ETs and in CX3cr1^gfp/gfp^ retinas, where macrophage/monocyte recruitment was impaired (**Fig. 4 a-b**). These findings suggest that macrophages/monocytes play a critical role in retinal repair through ETs. Inhibiting their recruitment or the release of ETs accelerated the recovery of the retinal parenchyma compared to broader neutrophil/macrophage depletion. Moreover, fluorescein angiography revealed the accumulation of sub-retinal fluid at day 1, with no statistical difference from the baseline in PAD4-treated mice, consistent with the absence of macrophage/monocyte infiltration (**Fig. 4 c-d**). The fluorescein leakage reduced similarly between the groups starting from day 4 (**Fig. 4 d**), but only the absence of ETs resulted in a gradual disappearance of the signal (**Fig. 4 c-d**). These results suggest that the blocking the generation of ETs is a more effective strategy for promoting retinal repair compared to blocking the recruitment of innate immunity into the retina.

**Fig. 4:**
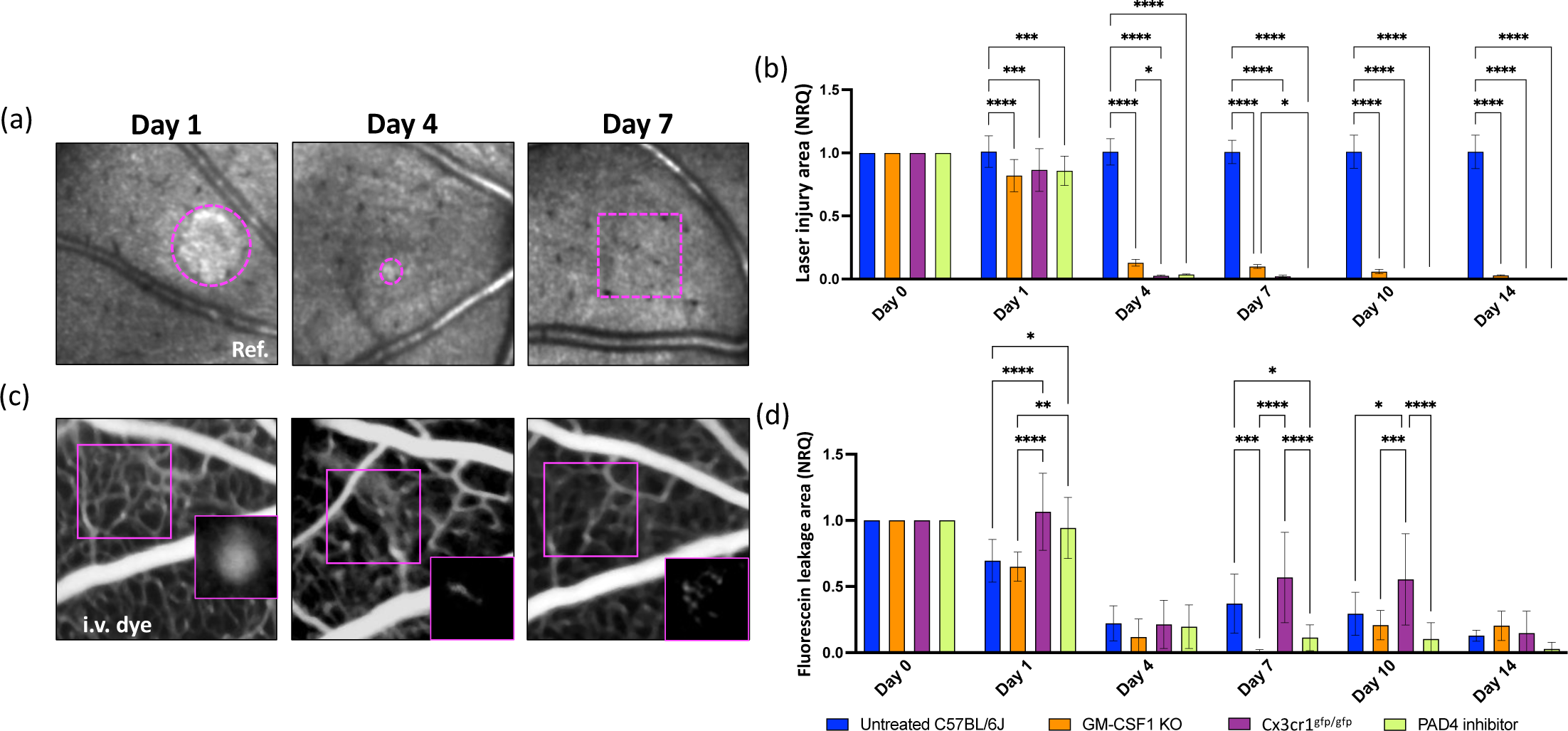
Impact of ETs on retinal and vascular repair upon laser-induced injury. (a-b) Kinetics of retinal injury detected in reflectance (back-scattered light) in untreated, GM-CSF1 KO, Cx3cr1^gfp/gfp^ and PAD4-treated mice. (a) Image of the same area after laser on days 1, 4 and 7 in PAD4-treated mice. We detected a reduction of hypo-/hyper-reflectivity in the damaged site on day 4, and no signal was found by day 7. (b) Quantification of the damaged area of untreated, GM-CSF1 KO, Cx3cr1^gfp/gfp^ and PAD4-treated retinas after injury (day 0) and at pre-defined time points (day 1, 4, 7, 10 and 14). Significant differences (*p<0.1, **p<0.01 and ****p<0.0001) between untreated, GM-CSF1 KO, Cx3cr1^gfp/gfp^ and PAD4-treated mice were determined by using a post-hoc Bonferroni one-way ANOVA test (n=8). (c) Representative fluorescein angiographs of the same PAD4-treated eyes on days 1, 4 and 7. (d) Quantification of dye that leaks in the depth of the retina, identified as leakage area, after injury (day 0) and at pre-defined time points (day 1, 4, 7, 10 and 14). Significant differences (*p<0.1, **p<0.01. ***p<0.001 and ****p<0.0001) between untreated, GM-CSF1 KO, Cx3cr1^gfp/gfp^ and PAD4-treated mice were determined by using a post-hoc Bonferroni one-way ANOVA test (n=8). For both groups, day 0 was chosen as calibrator [NRQ (normalized relative quantification) = 1]. Field of view is approximately ≈425 µm.

### ETs orchestrate the inflammatory response to laser-induced injury in the retina

In addition to known bactericidal and anti-fungal functions of ETs, recent studies also linked ET formation to sterile inflammation upon tissue damage (46). Indeed, ETs attract T cells and neutrophils that further amplify the pro-inflammatory response (47). To further establish the role of ETs during retinal inflammation, we pharmacologically inhibited ETs and analyzed *in vivo* how ETs affect the time course of PL and microglia response to injury in the retina and compared their inflammatory response to untreated mice (**Fig. 5 a-e**). PAD4 inhibition using Cl-amidine resulted in a quicker resolution of the inflammatory response (**Fig. 5 a, c, d**). Starting from day 1, microglia and PL and PL are recruited to the lesion similarly between PAD4-treated and untreated animals (**Fig. 5 c-d**). However, the density of GFP^+^ cells significantly diminished on day 4 and a slight difference was found compared to baseline (*p*=0.016; **Fig. 5 c**). Microglia accumulation in the lesion returned to baseline levels by day 7 only in PAD4-treated mice (**Fig. 5 a, c**). In both groups, PL were recruited to the injury starting from day 1, but the number of RFP^+^ cells in the injury was comparable to baseline by day 4 only after PAD treatment (**Fig. 5 a, d**).

**Fig. 5:**
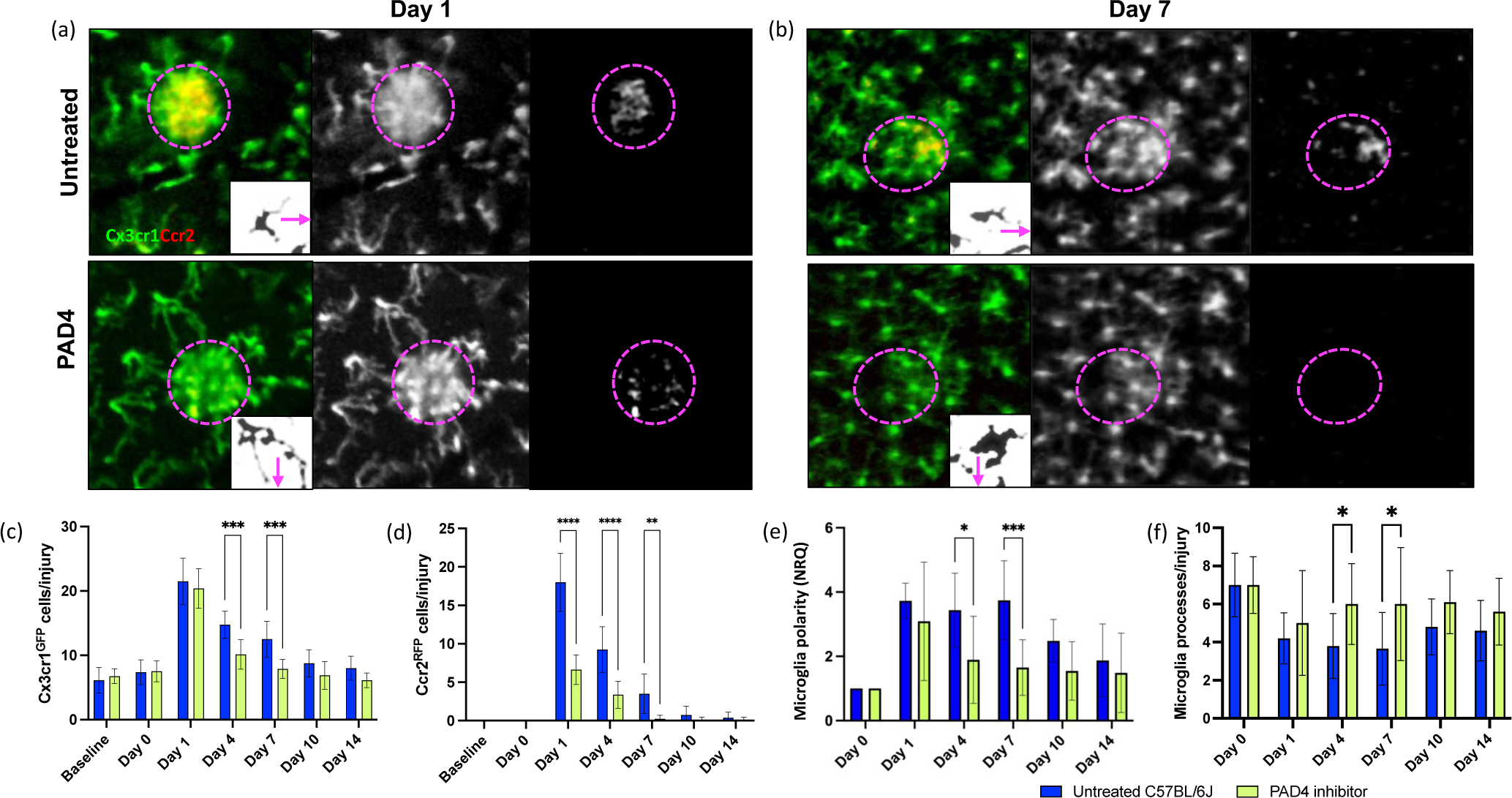
Repeated PAD4 dose regimen reduces retinal inflammatory upon laser-induced injury. (a-e) Inflammatory response of untreated and PAD4-treated Ccr2^RFP^Cx3cr1^GFP^ mice. (a-b) Images show differences in microglia and PL recruitment on days 1 and 7 in the damaged site (delimited by magenta dashes). While microglia and PL cluster in injury in untreated mice, PAD4-treated retinas have non-significant PL infiltration, and microglia are evenly distributed within the retinal parenchyma. Inserts show morphological stages of microglial activation in untreated and PAD4-treated mice. The arrow indicates the direction in which the damaged site is located. (c-d) Quantification of the number of GFP^+^ microglia and RFP^+^ PL per lesion of untreated and PAD4-treated Ccr2^RFP^Cx3cr1^GFP^ retinas before injury (baseline) and at pre-defined time points (day 0, 1, 4, 7, 10 and 14). Significant differences (**p<0.01, ***p<0.001 and ****p<0.0001) between untreated and PAD4-treated mice were determined by using a post-hoc Bonferroni one-way ANOVA test (n=8). (e) Quantification of the polarity coefficient of untreated and PAD4-treated microglia after injury (day 0) and at pre-defined time points (day 1, 4, 7, 10 and 14). Significant differences (*p<0.1 and ***p<0.001) between untreated and PAD4-treated mice were determined by using a post-hoc Bonferroni one-way ANOVA test (n=8). For both groups, day 0 was chosen as calibrator [NRQ (normalized relative quantification) = 1]. (f) Quantification of the primary and terminal processes per cell after injury (day 0) and at pre-defined time points (days 1, 4, 7, 10 and 14). Significant differences (*p<0. 1) between untreated and PAD4-treated mice were determined by using a post-hoc Bonferroni one-way ANOVA test (n=8). Field of view is approximately ≈425 µm.

Microglia display phenotypic plasticity in response to insult, reflecting their activation status. We assessed morphological changes and process orientation by measuring the polarization coefficient and process count per GFP^+^ microglia (**Fig. 5 b, e**). Before injury, both untreated and PAD4-treated mice exhibited quiescent microglia with ramified processes and no specific direction. However, upon injury, microglia became active and assumed an activated phenotype. Untreated microglia had fewer processes, but increased in length and orientation towards the injured site, indicating a pro-inflammatory state (**Fig. 5 b, e**). Similarly, PAD4-treated microglia showed polarization towards the injury at 24 hours post-injury, while between days 4 and 7, only PAD4-treated microglia demonstrated a rounded macrophage-like morphology with larger cell bodies and more processes, suggesting phagocytic activity during this time frame (**Fig. 5 b, e**). These findings indicate that ETs coordinate the inflammatory response, and the decrease in microglia and PL recruitment coincides with the morphological transformation of microglia towards a phagocytic phenotype.

We conducted an *in vivo* analysis to investigate in mice in which PAD4 expression was inhibited if and how ETosis control the inflammatory response to retinal injury (**Fig. 6 a-g**). We injected low doses of fluorescently labeled antibodies to differentiate neutrophils (Ly6G^+^), helper T-cells (CD4^+^), and cytotoxic T-cells (CD8^+^). We observed that CD4^+^ cells migrated toward the injury site within 24 hours and their density remained elevated for 4 days, coinciding with the clustering of CD8^+^ cells in the damaged area (**Fig. 6 b-c**). In contrast, the recruitment of neutrophils to the retina in response to injury was minimal (1-2 cells; Fig. 6 a, c). However, a significant number of Ly6G^+^ and CD8^+^ cells clustered at the superficial capillary plexus in the proximity of the injury site (Fig. 6 a, b, d). While a few neutrophils (∼2 cells) were consistently present above the injury, cytotoxic T-cells migrated closer to the damaged PR, particularly on day 4 (**Fig. 6 a, b, d**). We further examined the relationship between macrophages/monocytes and neutrophils, helper T-cells, and cytotoxic T-cells by correlating the number of Ly6G^+^, CD4^+^, and CD8^+^ cells with LysM^GFP^ cells clustering in the injured area of untreated mice (**Fig. S5**). We discovered a negative association between macrophages/monocytes and helper T-cells, but an increase in macrophages/monocytes occurred when Ly6G^+^ and CD8^+^ cells infiltrated during the injury response. These findings confirmed complex relation between macrophages/monocytes and these immune cell types, revealing both negative and positive associations, as shown by *in vivo* imaging after PAD4 treatment.

**Fig. 6:**
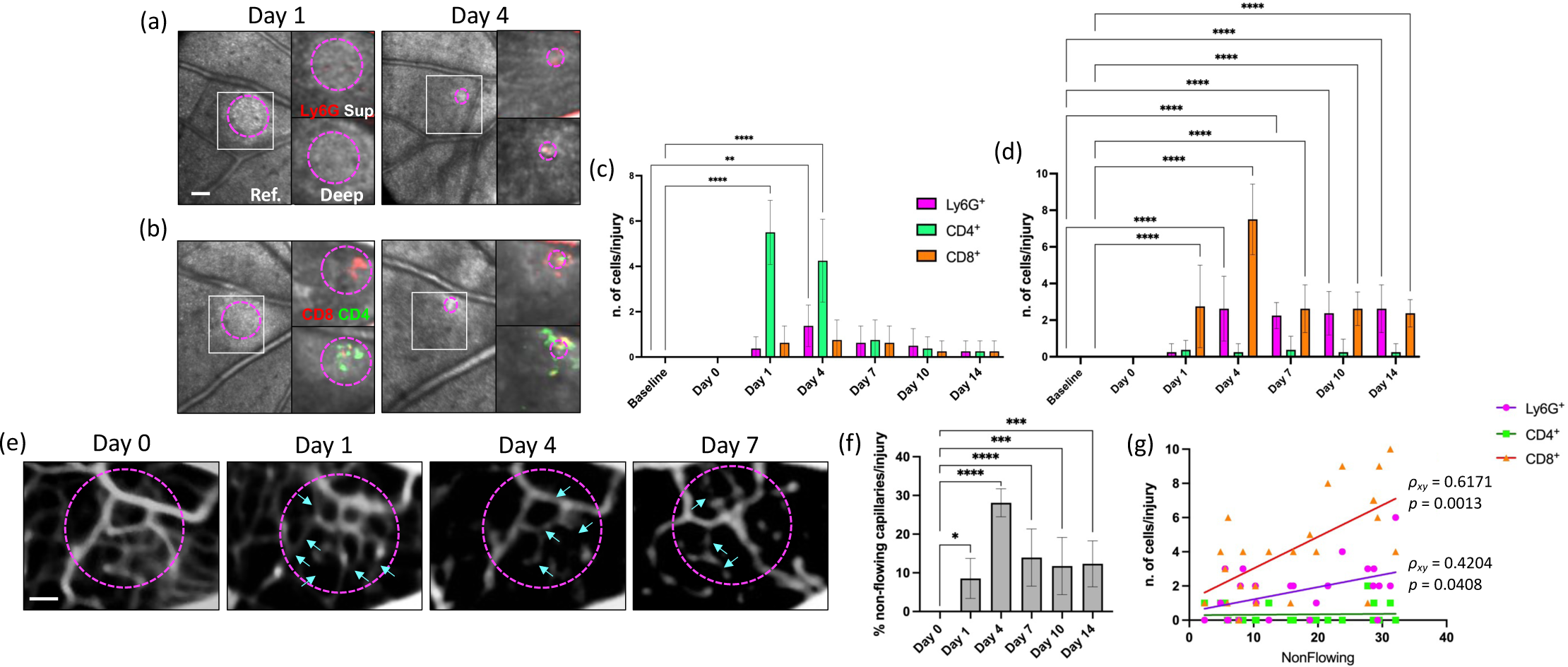
ETs orchestrate the inflammatory response to retinal injury. (a) Tracking of neutrophils (Ly6G^+^) in the same eye of a PAD4-treated mouse during injury response. Persistent number of CD4^+^ and CD8^+^ cells clustered in the damaged site. (b) Tracking of CD4^+^ and CD8^+^ T-cells in the same eye of a C57BL/6 mouse within the damaged area (magenta circles). The density of CD4^+^T-cells remained elevated for 4 days, when CD8^+^ cells also clustered in the damaged area. (c-d) Quantification of the number of neutrophils, CD4 and CD8 T-cells per lesion after laser in the proximity of the NFL (superficial/intermediate VP) and more in-depth by the PR at baseline and at different time points (Da 1, 4, 7, 10 and 14). Significant differences (***p<0.001 and ****p<0.0001) between baseline and the different time points were determined by using a post-hoc Bonferroni one-way ANOVA test (n=8). Field of view is approximately ≈575 µm. (e) Images of the anatomy and flow patterns of the superficial microvasculature obtained from the angiogram of PAD4-treated mice after injury (Day 0) and at different time points (Da 1, 4, 7, 10 and 14). We showed non-flowing capillaries with plugs (arrows) seen as filling defects, compared to day 1 when capillaries were flowing. (f) Percentage of non-flowing superficial capillaries per lesion after laser analyzed at different time points (Da 1, 4, 7, 10 and 14). Significant differences (*p<0.1, ***p<0.001 and ****p<0.0001) between baseline and the different time points were determined by using a post-hoc Bonferroni one-way ANOVA test (n=8). (g) Spearman correlation between capillary plugs per lesion with the number of neutrophils (Ly6G^+^), helper (CD4^+^), and cytotoxic T-cells (CD8^+^) clustering in the damaged site. Combined data from Days 4 and 7 after laser-injury.

We finally determined whether PL clustering at the level of the superficial capillary plexus results in plugging in the vasculature, leading to functional changes in retinal perfusion. We analyzed the anatomy and flow patterns of the microvasculature in the angiogram of PAD4-treated mice (**Fig. 6 e-g**). Starting from day 1, 8.5% of retinal capillaries in the proximity of the lesion exhibited no flow. By day 4, the proportion of plugged capillaries increased to 28.1% but returned to a level comparable to day 1 between days 7 and 14 (**Fig. 6 e-f**). We also discovered a positive correlation between non-flowing capillaries and the presence of neutrophils, helper T-cells, or cytotoxic T-cells at each time point analyzed (days 1-7; **Fig. S6**), suggesting that capillary plugging resulted from cellular blockage. Moreover, the severity of capillary plugging was higher when a more significant number of cytotoxic T-cells migrated closer to the damaged area (ρ_xy_= 0.6171). A weaker correlation was observed between non-flowing capillaries and neutrophils or helper T-cells (**Fig. 6 g**). These data suggest that PL clustering at the superficial capillary plexus leads to vasculature plugging, resulting in altered retinal perfusion. The temporal pattern of capillary obstruction, with a peak at day 4 and subsequent resolution, was linked to the presence of immune cells, particularly cytotoxic T-cells, highlighting their role in this process.

The absence of ETosis prevents PL from entering the damaged parenchyma; instead, they cluster in the superficial vasculature. This observation suggests that ETs play a role in regulating the immune response to laser-induced injury.

## DISCUSSION

Blood-borne macrophages/monocytes are the most prominent cells associated with chronic inflammation in degenerated retinas, outnumbering resident myeloid cells as well as lymphocytes (48, 49). Each macrophage/monocyte can secrete more than 100 molecules to alert other immune cells of diverse pathologic threats, thereby initiating and regulating multipronged immune responses (50). To date, the mechanisms by which macrophages/monocytes coordinate the dynamic inflammatory response locally and activation mechanisms remain unclear. Recent strides in experimental systems have allowed for real-time exploration of the intricate interactions among immune cells with distinct lineages and functionalities (51–53). Furthermore, these innovations have granted insights into the temporal dimensions of the response phase, encompassing its duration and timing (54). We used *in vivo* microscopy, electron microscopy, genetically modified mice to alter the innate immune response, and pharmacological treatments in a preclinical mouse model of retinal degeneration to study how macrophages/monocytes influence the spatio-temporal dynamics of microglia and PL responses.

Through TEM analysis, we have observed that phagocytic cells release chromatin and granular proteins, forming ETs that play a pivotal role in tissue repair. In contrast to expectations, Neutrophils did not produce ETs in this context. Combining RNA-seq and *in vivo* imaging confirmed that macrophages/monocytes are capable of undergoing ETosis. Blocking this process using Cl-amidine promoted more effective retinal and vasculature repair and hindered the recruitment of innate immune cells into the injured parenchyma. Cl-amidine indeed induced alterations in the inflammatory response, leading to the clustering of neutrophils, helper T-cells, and cytotoxic T-cells primarily at the level of the superficial capillary plexus, causing capillary obstruction. Our data demonstrate that macrophages/monocytes play a crucial role in coordinating the immune response to laser-induced injury by forming ETs.

Despite various clinical trials aiming to target inflammatory pathways within the retina, their outcomes have been largely unsuccessful, and the reasons behind their failures remain unclear (55). A limiting factor is the relative insufficient knowledge of the specific role of the different immune cell types within the retinal inflammatory orchestra that leads to tissue degeneration. Genetic investigations (56), alongside human histological data and animal studies (57, 58), have predominantly concentrated on the involvement of innate immunity in mediating cellular death during the progression of retinal degeneration. Upon injury, macrophages/monocytes are activated, and their reactivity can be harmful as well as beneficial, depending on the surrounding milieu. Initially, they adapt the classically activated state that improves their phagocytic function. However, macrophages/monocytes can also assume an alternatively activated phenotype during prolonged inflammation that fuel the inflammatory process. Similarly, neutrophils also wield the ability to stimulate both acute and chronic inflammation. They are the first responders to acute inflammation contributing to the resolution of inflammation. However, during chronic inflammation, they can cause tissue damage and intensify the immune response. Nevertheless, the extent to which macrophages/monocytes and neutrophils contribute to retinal degeneration remains controversial.

Here, we provide evidence of the involvement of innate immunity during laser-induced injury response in mice. The laser-induced injury model has shown to be relevant for studying human neurodegeneration in the retina as it mimics certain macroscopic features of human retinal degenerative diseases (59). Moreover, our injury model is an excellent resource for investigating specific biological processes directly in the area of damage. In support of this, previous studies have provided *in vivo* evidence that resident and blood-borne immune cells migrate into the area of injury in the retina following the induction of light-induced retinal lesions (24, 60). We found that innate immune cells were recruited close to the damaged area on both days 1 and 7, with a reduction in their response on day 4. Macrophages/monocytes clustered above the damage site on days 1 and 7 and were primarily present around the injured PR on day 1. Contrariwise, neutrophils appeared near the damaged outer retina on days 1 and 7 and migrated towards the injured site in the proximity of the NFL on mostly day 1. These results imply that macrophages/monocytes and neutrophils migrated to the injured area in two waves with distinct patterns of cellular migration and temporal changes. Our results are in line with the literature that shows early recruitment of innate immune cells, followed by a secondary cell infiltration from the bloodstream, which is mediated by both tissue-resident and early-recruited cells (61, 62). Interestingly, the activation status affected their migratory patterns (63), implying that macrophages/monocytes and neutrophils migrated to the injury on day 1 or 7 may have different effects on tissue repair in the retina.

We then analyzed the impact of innate immunity on retinal repair using a mouse model unable to generate neutrophils and macrophages, GM-CSF1 KO mice, and Cx3cr1^gfp/gfp^ mice, which exhibit impaired recruitment of macrophages/monocytes. GM-CSF is a hematopoietic growth factor controlling mature myeloid cell populations under homeostatic and inflammatory conditions (64, 65). It supports the activation of mature neutrophils and macrophages, inducing the production of proinflammatory cytokines and enhancing their survival and activation (66, 67). Cx3CR1, or fractalkine receptor, coordinates the recruitment of macrophages/monocytes to sites of damage and is a key regulator of their function at sites of inflammation. (68, 69). In several murine models of autoimmunity/inflammation, GM-CSF blockade led to reduced levels of monocyte and neutrophil recruitment with corresponding alleviation of disease severity (70–72). While homozygous mutant Cx3cr1^gfp/gfp^ mice lacking CX3CR1 expression (22) were shown to have a selective anti-inflammatory effect promoting neuroprotection by reducing the recruitment of macrophages/monocytes (73). Similarly, we showed *in vivo* the crucial role of innate immunity in retinal repair, as the extent of the damage reduced 24 hours post injury in both mutant mice compared to wild-type (WT) animals. Remarkably, the impaired recruitment of macrophages/monocytes accelerated further the recovery of the retinal parenchyma compared to the broader depletion of neutrophils and macrophages. The different outcome obtained from those mutant mice can be due to the presence of neutrophils in Cx3cr1^gfp/gfp^ mice as fractalkine is not expressed on neutrophils nor stimulate migration of these cells. Neutrophils are known to lead to tissue damage (74); however, they can exert neuroprotection in the central nervous system (CNS) and play critical roles in antiinflammation (75, 76). For instance, neutrophils secrete several molecules that are beneficial for promoting neuron cell survival, such as granulocyte-colony stimulating factor (G-CSF), hepatocyte growth factor (HGF), nerve growth factor (NGF), and neurotrophin-4 (NT4) (77–81).

Concurrently, we analyzed *in vivo* the integrity of the retinal vasculature during injury response between GM-CSF1 KO, Cx3cr1^gfp/gfp^ and WT animals. The vascular network adjacent to the PR was compromised until the last time point investigated in all groups; however, untreated and Cx3cr1^gfp/gfp^ mice exhibited a more pronounced leakage between days 7 and 10 compared to animals lacking of macrophage and neutrophil recruitment. Hence, the depletion of not only macrophages/monocytes but predominantly neutrophils could prove advantageous in restoring vascular integrity. Neutrophils, in fact, break blood barriers significantly due to their abnormal interactions with the endothelium (82). Their capacity to degrade neurovascular units following injury, mediated by matrix metalloproteinases (e.g., MMP2, MMP3, and MMP9) and reactive agents like reactive oxygen species (ROS), nitrogen oxides (NOS), and NADPH oxidase, underscores their impact (83, 84). Particularly, these molecular species induce direct oxidative harm and reorganization of tight junctions, contributing substantially to the disruption of blood barrier integrity (85, 86). The statistical analysis of our data solely linked the accumulation of macrophages/monocytes at the injured site, rather than neutrophils, to the extent of parenchymal and vascular damage. This finding corroborates that a targeted depletion of macrophages/monocytes among the innate immune cell population could present an efficacious approach for facilitating retinal recovery. Nevertheless, it is challenging to address whether the beneficial impact of Cx3cr1 deletion on retinal repair is due to its impact on macrophage/monocyte recruitment or a direct effect on microglial activation and survival (87, 88). These results suggest that a targeted depletion of macrophages/monocytes among the innate immune cell population could present an efficacious approach for facilitating retinal recovery. Nevertheless, it is challenging to address whether the beneficial impact of Cx3cr1 deletion on retinal repair is due to its impact on macrophage/monocyte recruitment or a direct effect on microglial activation (87, 88). Previous research attempted to study the role of macrophages and microglia separately; however, it has been challenging. Once macrophages/monocytes infiltrate the retina and cluster in the injured area, they become virtually indistinguishable from resident microglia not only morphologically but also in surface markers and function (89). These similarities have complicated the development of efficient treatments to deplete one or the other cell types. Different major pharmacological approaches are now in use to investigate macrophage and microglia functions during CNS injury response such as CSF1R inhibitors (e.g., PLX5622 and PLX3397). CSF1R inhibitors have been suggested to selectively deplete CNS microglia without a significant impact on peripheral immune cells (90–92). Nevertheless, their influence extends beyond microglia, encompassing lasting alterations within the myeloid populations of the bone marrow, spleen, and bloodstream (93). For instance, PLX compounds suppress the proliferation of monocyte progenitor cells and macrophages derived from the bone marrow (94, 95). Furthermore, these inhibitors hamper the functionality of macrophages in the bone marrow and spleen, as evidenced by a diminished phagocytosis (93). Previous studies have shown that microglia depletion by CSF1R inhibition can either promote or exacerbate neurodegeneration (96, 97). These conflicting results might arise not solely from the divergent functions of microglia across distinct CNS disease models but also from the various contribution of peripheral and circulating macrophages. Recognizing the intricate role of innate immunity in CNS repair, we focused on enhancing their capacity to effectively orchestrate the immune response, rather than delving into alternative depletion mechanisms.

Innate immune cells can release extracellular structures known as ETs (98). Apart from their recognized bactericidal and anti-fungal functions (99, 100), recent research has unveiled a connection between ET formation and sterile inflammation following tissue damage (101). Domer et al. demonstrated how ETs could induce further ET formation and establish a positive feedback loop capable of intensifying inflammation (102). Consequently, the persistent ETosis offers the basis for driving persistent inflammatory responses. Additionally, ET release have been documented in preclinical mouse models of ocular inflammation, as well as in samples collected through standard pars plana vitrectomy from diabetic patients (14). ETosis comprises a unique series of cellular events by which nuclear contents, including chromatin, mix with cellular proteins and are then extruded from the cell body to form extracellular structures capable of “trapping” and killing microorganisms. Since the original report about neutrophils, other leukocytes including macrophages/monocytes are now known to produce ET structures. In our injury model, neutrophils displayed a characteristic thin cell membrane, signifying that they were not undergoing ETosis after injury. On the other hand, phagocytic cells exhibited disintegrated nuclear membranes and decondensed chromatin. Chromatin either accumulated within cytoplasmic blebs or was released into the extracellular space, typical of ETosis (35). Furthermore, our investigation identified a significant up-regulation of 38 genes related to ETosis in phagocytic cells (Csf1r^+^) during the injury response, confirming their ability of releasing ETs. Both microglia and macrophages were found to participate in ET formation (103, 104), suggesting ETosis as an alternative defense mechanism when cellular phagocytic capacity becomes overwhelmed (105). We employed *in vivo* confocal imaging using a DNA-binding dye to visualize ET formation. This allowed us to quantify the release of ETs by microglia (Cx3cr1^+^) and macrophages (LysM^GFP^) in response to laser-induced retinal injury in reporter mice. Interestingly, macrophages exhibited a preference for ETosis, as indicated by the preferential co-localization of the DNA dye SYTOX with LysM^GFP^ cells.

Histone citrullination is the prerequisite and trigger for ET formation, which is mediated by PADs via converting arginine to citrulline (106, 107). The citrullination level of histones, chromatin decondensation, and ETs-like structure formation are susceptible to PAD4 activity (108). In line with the literature, our transcriptome analysis of phagocytic cells revealed an increased expression of PAD4. The PAD4 inhibitors in preclinical models of CNS injury, such as choroidal neovascularization and traumatic brain injury (TBI), demonstrated that the level of ETs formation *in vivo* is closely correlated with local inflammation and pathogenesis (109–111). Vaibhav et al. recently reported that the administration of Cl-amidine led to reduced ET production post-TBI, accompanied by decreased cerebral edema, improved blood flow, and alleviated neurological deficits following the injury (111). Our lab also proved that Cl-amidine resulted in a quicker resolution of the inflammatory response. Cx3cr1^+^ cells returned to baseline levels by day 7 only after treatment. However, the reduction was clearly visible on day 4, concomitant with the absence of PL clustering in the injured site. The role of ETs in host defense complements phagocytosis (112). As reported in literature, we observed *in vivo* that microglia exhibit a rounded macrophage-like morphology, indicating phagocytic activity only when ETosis is inhibited (113). Conversely, untreated microglia extended processes towards the injured site, signifying a pro-inflammatory state (114). Thus, ETosis appears to act as an alternative mechanism when phagocytosis is hindered, potentially exacerbating the inflammatory response. In support of these latter points, recent studies demonstrated that phagocytosis and ETosis represent alternative outcomes of innate immune cell activation, which may adjust negatively to each other (115). However, signals that determine the choice between phagocytosis and the generation of ETs are still poorly characterized.

Nevertheless, it is challenging to address if the beneficial impact of ET inhibition on retinal repair is due to its effect on the overall inflammatory response or a cellular response of one contributor that infiltrated the retina. To address this, we analyzed separately different sub-populations of PL known to exert detrimental effects on retinal repair. Following the inhibition of ETosis, we observed distinct migration patterns in neutrophils and T-cells. Neutrophils and cytotoxic T-cells clustered at the injury site within 24 hours of its induction, maintaining a consistent presence above it. Cytotoxic T-cells gradually migrated closer to the damaged photoreceptors, particularly by day 4. Additionally, helper T-cells exhibited rapid migration toward the injury site within the first 24 hours, with their density remaining elevated for 4 days. Although correlation merely explains how strong the relationship between quantitative variables may be and not the causality between them, we observed a positive correlation between the recruitment of macrophages/monocytes and the infiltration of neutrophils and cytotoxic T-cells during the injury response. Conversely, there was a negative correlation with helper T-cells. Consequently, macrophages/monocytes’ engagement in the injury response and their ability to release ETs may emerge as critical factors contributing to the infiltration of neutrophils and T-cells. Remarkably, little is known about the effect of ETs on neutrophils during injury response. Neutrophils play a pivotal role as the initial responders to inflammation, often constituting the predominant leukocyte population at the inflammatory site (116, 117). Consequently, they are likely the first immune cells to encounter ETs released during the early stages of the inflammatory response. Therefore, activation of neutrophils by ETs has the potential to exert a strong regulatory effect on the progression of inflammation (102). ETs are also known to activate various effector functions within neutrophils, including exocytosis, the generation of ROS, the formation of their own ETs, as well as the process of phagocytosis (118). Furthermore, NETs can stimulate the release of proinflammatory chemokines, further intensifying the inflammatory milieu. ETs also play a significant role in activating T-cells through their histones, establishing a novel link between innate and adaptive immune responses (119). ETs can prime T-cells by lowering their activation threshold and enhancing T-cell responses to specific antigens (120). However, this effect isn’t equal across all T-cell subpopulations (121–123). We recently showed the involvement of adaptive immunity in retinal damage (29), and combining our data with these previous findings, we can conclude that T-cell responses is partially altered by ET inhibition. The recruitment of helper T-cells by the injured PR was preserved after Cl-amidine, while ET inhibition hinders CD8^+^ cell response to injury. This appeared as a reduction in the number of CD8^+^ cells reaching the site of injury. Instead of accumulating within the retinal parenchyma on day 4 post-injury, cytotoxic T-cells were predominantly observed near the superficial capillary plexus, just above the damaged area. Although neutrophils usually considered as the first responders, macrophages/monocytes are recruited within the first few hours of injury (124). In support of our data, macrophages/monocytes and neutrophils are known to collaborate in various disease contexts, and their intercellular communication plays a pivotal role in shaping the outcomes of tissue repair (124). Once activated in response to tissue damage, macrophages attract neutrophils to the inflammatory site by secreting chemoattractant such as CXCL1, CXCL2, IL1α, and CCL2. Additionally, they extend the lifespan of neutrophils by releasing growth factors and cytokines like GM-CSF, G-CSF, and TNFα (125). Active neutrophils, in turn, further fuel the inflammatory response by recruiting additional monocytes and influencing their differentiation and macrophage polarization (126). While macrophages/monocytes initiate the inflammatory response and induce the release of various inflammatory cytokines, T-cells are recruited to the injured CNS and release pro-inflammatory and anti-inflammatory cytokines (127–129). Once those cell types reach the injury, their interplay maintains tissue homeostasis and orchestrates wound healing (130–132). Notably, T-cell differentiation and macrophage polarization exert precise control over the tissue microenvironment in response to injury, and their actions often intersect (133). For instance, certain subsets of T-cells can activate pro-inflammatory macrophages through the secretion of IFN-γ, while the production of IL4 by other T-cell subsets promotes the polarization of macrophages toward an anti-inflammatory phenotype (134). Conversely, macrophages contribute to the milieu by releasing pro-inflammatory chemokines and simultaneously inhibit the recruitment of T-cells (135).

Finally, we aimed to understand whether PL recruitment at the superficial capillary plexus can influence the perfusion of retinal microvasculature. Previous studies indicated that capillary obstruction could potentially contribute to CNS injury. Even a minimal obstruction of 1.8% of capillaries due to stalled PL has been shown to result in a reduction in overall cerebral blood flow, with a direct correlation observed between the fraction of plugged capillaries and neurological dysfunction (136). Our data are consistent with the idea that PL, particularly neutrophils and cytotoxic T-cells, have an impact on the microvascular blood flow when their migration patterns are altered by ETosis inhibition. In our focal and acute injury model, a decrease in blood flow was evident 24 hours after damage when ET inhibition was performed. This reduction coincided with an increase in the occurrence of plugged capillaries, reaching a peak of 28.1% by day 4. We also found a positive correlation between obstructed capillaries and the presence of neutrophils and T-cells. This correlation became more pronounced when a substantial number of cytotoxic T-cells migrated closer to the damaged area. These findings suggest that ETosis is crucial in guiding neutrophils and T-cells toward the injured PR. In its absence, PL stalls in the microvascular bed, contributing to increased flow resistance. Since the first description of ETs more than a decade ago (137), ETosis has been recognized as a significant aspect of damage in various CNS diseases (138, 139). Recent research has uncovered evidence of ETs and their involvement in the pathophysiology of ocular diseases, including diabetic retinopathy, uveitis, and AMD (14, 15, 140). Relevant to our findings, Chen et al. demonstrated that the amyloid β-protein Aβ1-40, the primary component of drusen, triggers the release of ETs through the activation of pro-inflammatory pathways (141). Their findings also demonstrated that PAD4 inhibitors effectively alleviate PL infiltration in the retina and, thus, chronic inflammation.

In conclusion, our data suggest a critical function of PAD4-mediated ETosis performed by macrophages/monocytes during retinal injury, indicating that targeting infiltrating phagocytic cells capable of releasing ETs may be a potential treatment for retinal degeneration. Novel inhibitors of the PAD4 activity, already in preclinical studies for non-ocular diseases, may also provide novel strategies to stop these early events associated with the development of retinal degeneration. Nevertheless, future studies are necessary to research ETosis in retinal degenerative diseases further. A more detailed characterization of these ETs-releasing phagocytic cells is also of interest to determine whether different phenotypes of macrophages/monocytes have distinct roles upon retinal injury (e.g., sc-RNAseq), which would further complement our results. Thus, understanding these pathological pathways may pave the way for new therapeutic opportunities for retinal degenerative diseases like AMD.

