## Supplementary figures for "MACROPHAGES COORDINATE IMMUNE RESPONSE TO LASER-INDUCED INJURY VIA EXTRACELLULAR TRAPS"

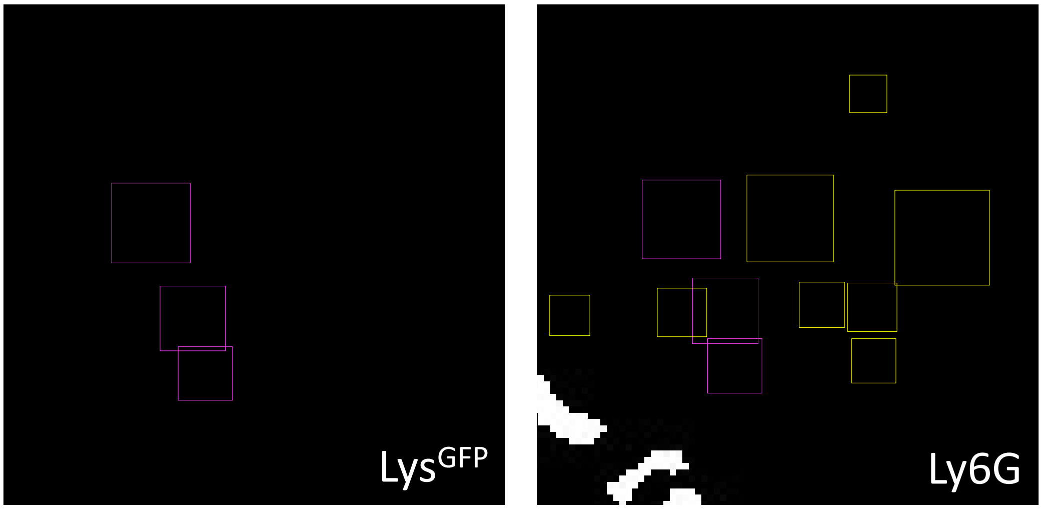


***Fig. S1: Image processing for the quantification of innate immune response in pictures obtained by in vivo imaging.*** *Representative image shows the approach for counting macrophages/monocytes (LysM$^{GFP}$) and neutrophils (LysM$^{GFP}$/Ly6G$^{+}$). It relies only on ImageJ thresholding. LysM$^{GFP}$ are surrounded by a yellow frame and LysM$^{GFP}$/Ly6G$^{+}$ cells are highlighted by a magenta frame.*

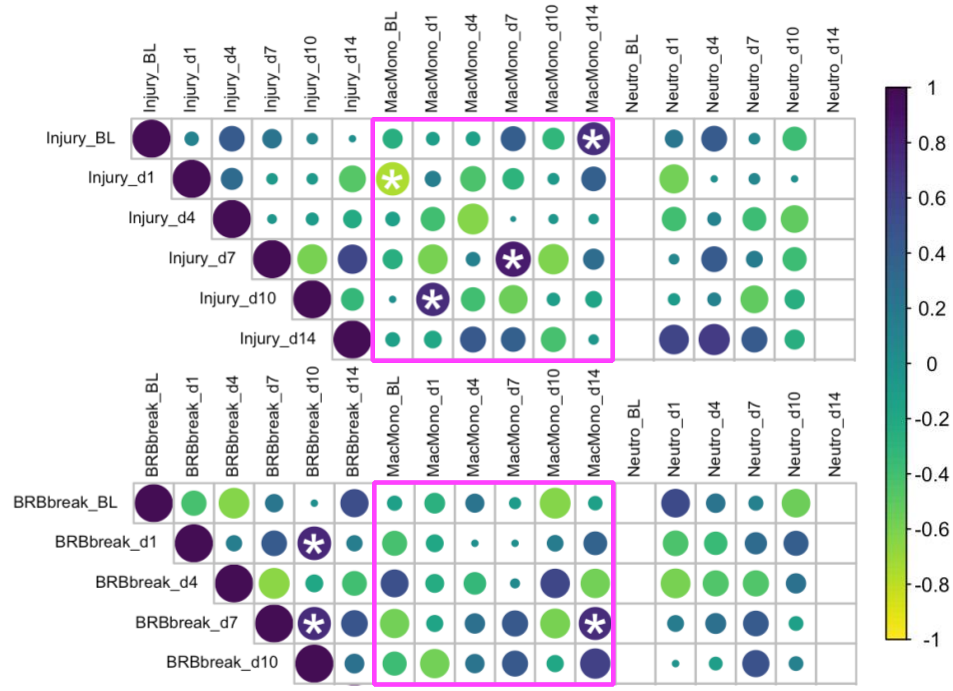


***Fig. S2: Association between the clustering of macrophages/monocytes with parenchymal and vascular damage.*** *Spearman's rank-order correlation between damaged (Injury, top) and the area of fluorescein leakage (BRB break, bottom) with the number of macrophages/monocytes (LysM$^{GFP}$) and neutrophils (LysM$^{GFP}$/Ly6G$^{+}$) clustering in the injury in untreated mice. Color intensity and the size of the circle are proportional to the correlation coefficients, and a star (*) marks significant correlation.*

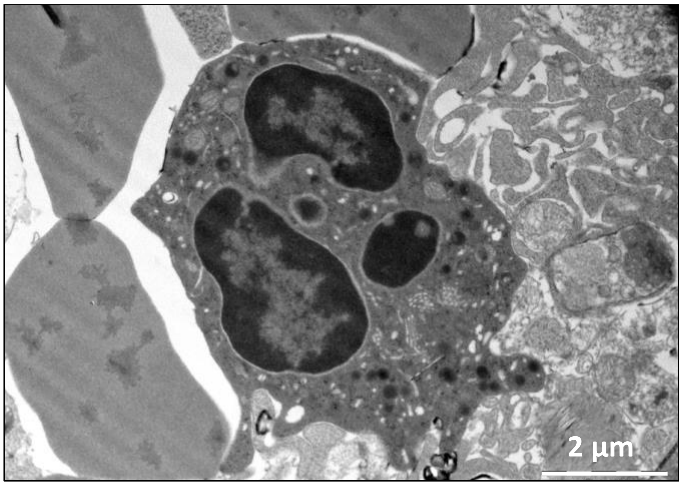


***Fig. S3: Absence of ETosis in neutrophils during injury response in the retina.*** *Transmission electron microscopy showing a neutrophil in the retinal tissue and no of ET formation was observed. This cell displays various granules in the cytoplasm, along with a lobulated nucleus. The condensed heterochromatin (dark) is positioned at the nucleus's edge, interrupted by euchromatic areas near nuclear pores. The brighter euchromatin is predominantly located in the center of the lobules.*

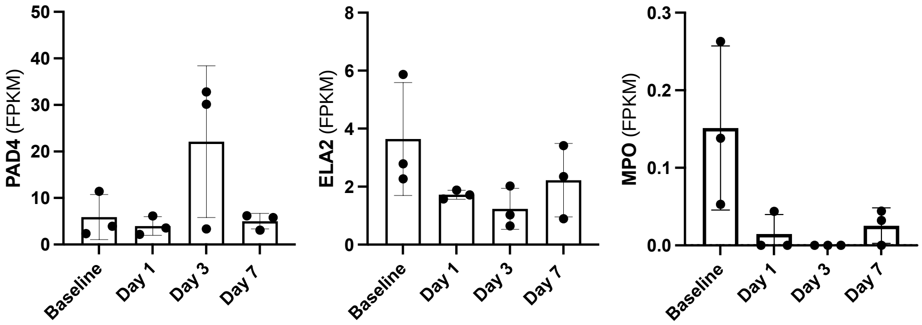


***Fig. S4: Gene expression of upstream regulators of ET formation.*** RNA-seq data illustrates the gene expression values of PAD4, ELA2, and MPO, expressed as fragments per kilobase of transcript per million mapped reads (FPKM).

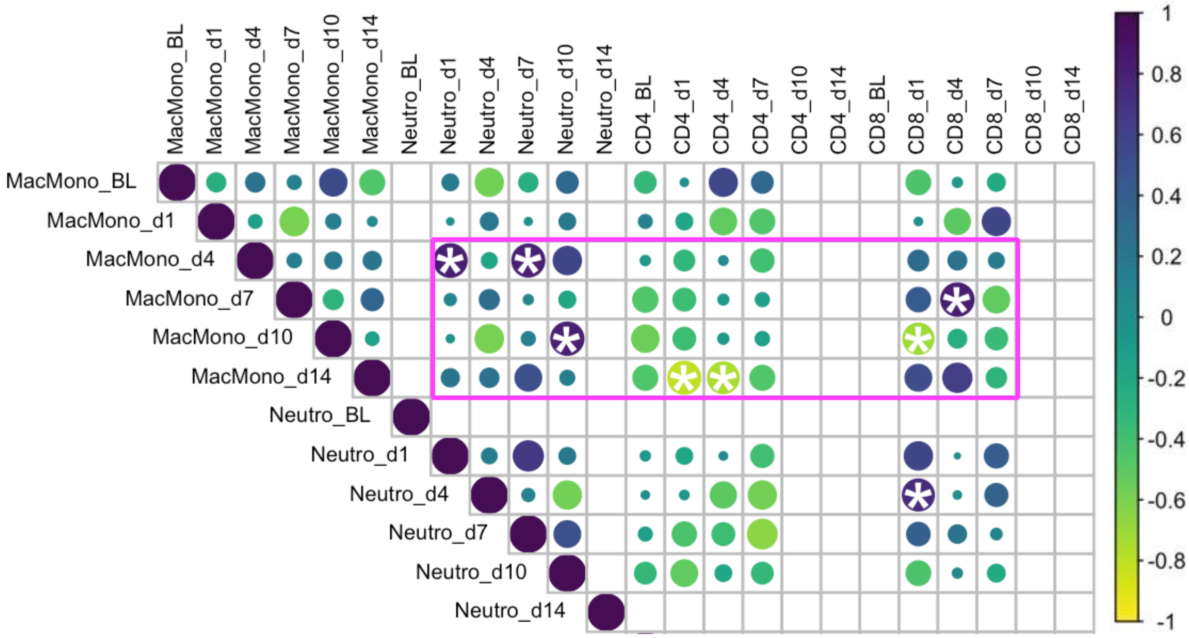

***Fig. S5: Association between the clustering of innate immune cells with neutrophils and T-cells.*** *Spearman's rank-order correlation between the number of macrophages/monocytes (LysM$^{GFP}$), neutrophils (LysM$^{GFP}$/Ly6G$^{+}$), CD4$^{+}$ and CD8$^{+}$ T-cells clustering in the injury. Color intensity and the size of the circle are proportional to the correlation coefficients, and a star (*) marks significant correlation.*

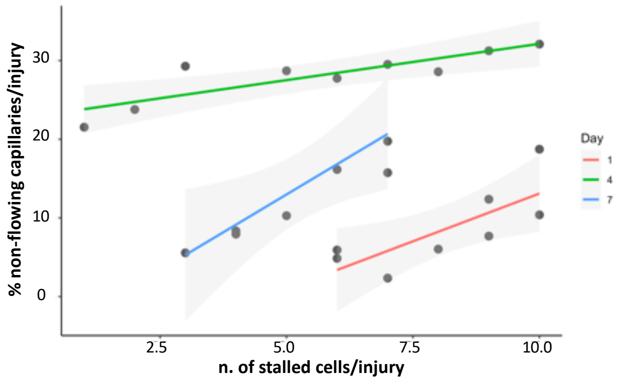


***Fig. S6: Association between the capillary plugs and stalled cells in the capillaries.*** *Spearman correlation between non-flowing capillaries with plugs with the number of cells stalled in the injured area nearby the NFL.*
